# Movement ecology of vulnerable lowland tapirs across a gradient of human disturbance

**DOI:** 10.1101/2021.11.12.468362

**Authors:** EP Medici, S Mezzini, CH Fleming, JM Calabrese, MJ Noonan

## Abstract

Animal movement is a key ecological process that is tightly coupled to local environmental conditions. While agriculture, urbanisation, and transportation infrastructure are critical to human socio-economic improvement, these have spurred substantial changes in animal movement across the globe with potential impacts on fitness and survival. Notably, however, human disturbance can have differential effects across species, and responses to human activities are thus largely taxa and context specific. As human disturbance is only expected to worsen over the next decade it is critical to better understand how species respond to human disturbance in order to develop effective, case-specific conservation strategies. Here, we use an extensive telemetry dataset collected over 22 years to fill a critical knowledge gap in the movement ecology of lowland tapirs (*Tapirus terrestris*) across a gradient of human disturbance within three biomes in southern Brazil: the Pantanal, Cerrado, and Atlantic Forest.

From these data we found that the mean home range size across all monitored tapirs was 8.31 km^2^ (95% CI: 6.53 - 10.42), with no evidence that home range sizes differed between sexes nor age groups. Interestingly, although the Atlantic Forest, Cerrado, and Pantanal vary substantially in habitat composition, levels of human disturbance, and tapir population densities, we found that lowland tapir movement behaviour and space use were consistent across all three biomes. Human disturbance also had no detectable effect on lowland tapir movement. Lowland tapirs living in the most altered habitats we monitored exhibited movement behaviour that was comparable to that of tapirs living in a near pristine environment.

Contrary to our expectations, we observed very little individual variability in lowland tapir space use and movement, and human impacts on the landscape also had no measurable effect on their movement. Lowland tapir movement behaviour thus appears to exhibit very little phenotypic plasticity. Crucially, the lack of any detectable response to anthropogenic disturbance suggests that human modified habitats risk being ecological traps for tapirs and this information should be factored into conservation actions and species management aimed towards protecting lowland tapir populations.

## Introduction

While agriculture, urbanisation, and transportation infrastructure are critical to human socio-economic improvement (Esfahani and Ramírez 2003), the associated habitat transformations represent a major threat to species survival (Fahrig 1997; Venter et al. 2006; Powers and Jetz 2019). Of particular concern is the impact of human activities on animal movement and space use (Allen and Singh 2016; Tucker et al. 2018; Doherty et al. 2021). Animal movement governs how individuals, populations, and species interact with each other and the environment (Schick et al. 2008; Martinez-Garcia et al. 2020; He et al. 2021) and mediates key ecological processes (Bauer and Hoye 2014). The capacity for individuals to move unhindered across complex landscapes is therefore critical for species survival and ecosystem function. Problematically, human development has reduced the amount of habitat available to wildlife (Brooks et al. 2002; Cardinale et al. 2012; Hooper et al. 2012). This has spurred substantial changes in animal movement behaviour across the globe (Fahrig 2007; Tucker et al. 2018; Doherty et al. 2021), with potential consequences including reduced fitness and survival, altered predator-prey dynamics, reduced seed dispersal, genetic isolation and local extinction (Fahrig 2007; Dickie et al. 2017; Cosgrove et al. 2018; Tucker et al. 2021).

Notably, human disturbance has been shown to have differential effects across species (Toews et al. 2018; Doherty et al. 2021), even for closely related taxa occupying the same habitat (Thatte et al. 2020). Responses to human activities are thus largely context specific (Doherty et al. 2021) and cannot be expected to be consistent across taxa. For instance, while Wall et al. (2021) found a tendency for African elephants (*Loxodonta spp*.) to exhibit reduced movement in human modified landscapes, Morato et al. (2016) noted that jaguars (*Panthera onca*) living in regions with high human population densities in South America occupied home ranges that were orders of magnitude larger than those of jaguars living in more pristine habitats. As human disturbance is only expected to worsen over the next decade it is critical to better understand how species respond to human disturbance to develop effective, case-specific conservation strategies.

Here we focus on understanding how the movement behaviour of lowland tapirs (*Tapirus terrestris*) varies across a gradient of human disturbance within the Pantanal, Cerrado, and Atlantic Forest biomes in southern Brazil. Lowland tapirs are herbivores of the order Perissodactyla that can reach over 2.5 meters in length and weigh up to 250kg (Medici 2011). While lowland tapirs are distributed throughout South America (Gardner 2008), their populations have suffered severe reductions, with local and regional extirpations, and are currently classified as vulnerable to extinction (Varela et al. 2019). Although the incorporation of information on animal movement is a key component in designing effective conservation and recovery strategies (Allen and Singh 2016), currently, very little is known about the movement ecology of tapirs (but see Noss et al. 2003; Tobler 2008; Fleming et al. 2019). This knowledge gap is especially pertinent given that large terrestrial mammals, such as tapirs, tend to have larger home ranges and greater absolute mobility than do small mammals (Calder 1983; Noonan et al. 2020), making them more susceptible to anthropogenic impacts than smaller bodied species (Tucker et al. 2018; Hill et al. 2020). Here, we use an extensive telemetry dataset collected over 22 years to describe the movement ecology of tapirs and study how changes in habitat composition and human disturbance influence their movement and space use. First, animals living in highly productive environments do not need to range over wide areas to meet their energetic needs (Lucherini and Lovari 1996; Relyea et al. 2000; Nilsen et al. 2005). As such, we expected that tapirs should exhibit plasticity in their movement and space use in relation to local environmental conditions as well as biome type. Furthermore, because human activity tends to result in increased movement for large herbivores (Doherty et al. 2021) our underlying hypothesis was that tapirs should exhibit greater movement distances and larger home range areas when living in human-modified landscapes.

## Methods

### Study area and data collection

The data were collected in three different biomes in southern Brazil (Fig. 1): Atlantic Forest (1997-2007), Pantanal (2008-2019), and south-western Cerrado (2016-2018).

**Figure 1:**
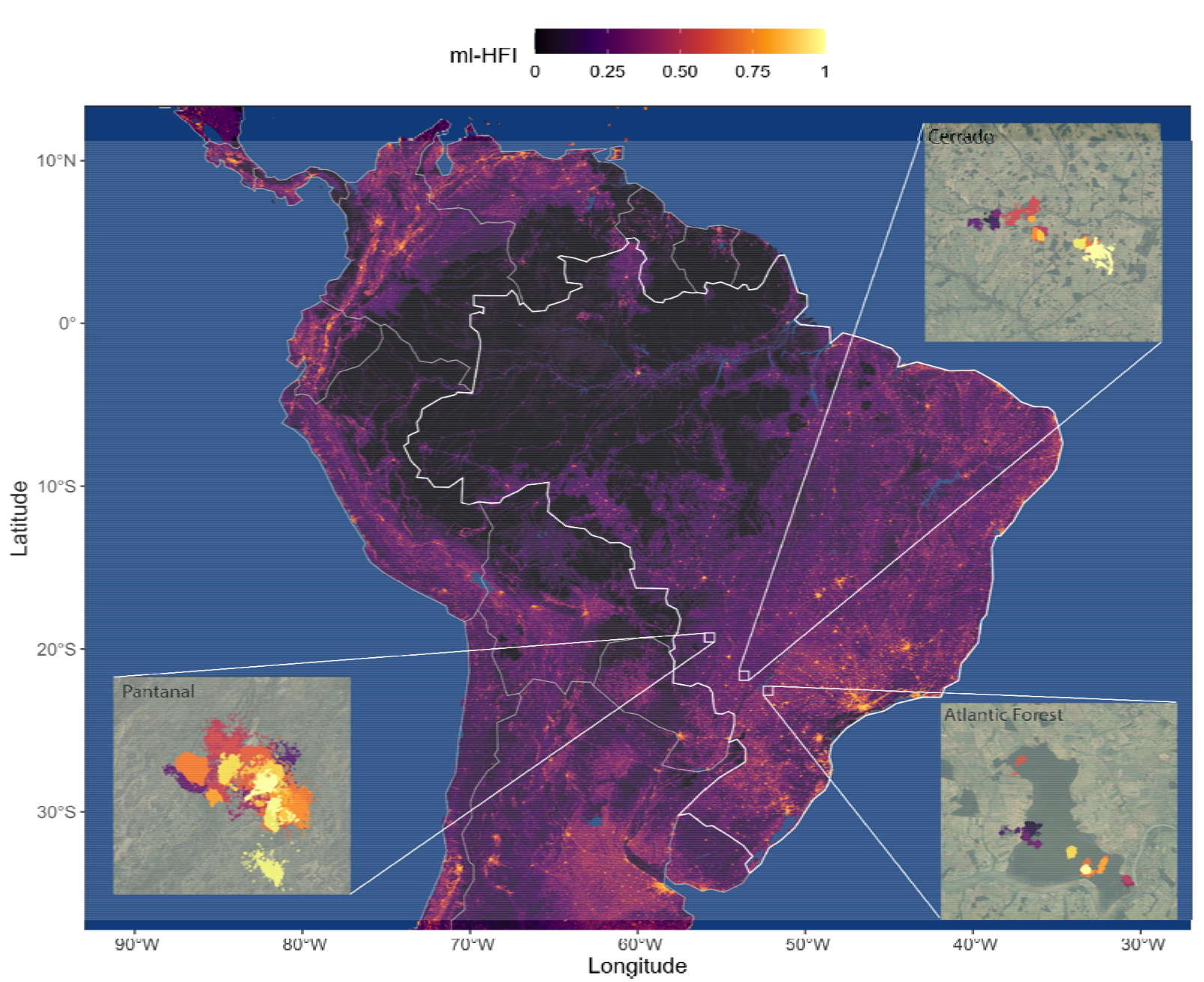
Location of the tree study sites (Pantanal, Cerrado, Atlantic Forest) over a raster of machine-learning-based human footprint index (ml-HFI), an index of human pressure on the landscape that is derived from remotely sensed surface imagery and ranges on a scale between 0 (no human impact), and 1 (high human impact).

#### Atlantic Forest

Morro do Diabo State Park is a protected area located in the Municipality of Teodoro Sampaio (22°32’S, 52°18’W), state of São Paulo, in the southeastern region of Brazil. The park has an area of 370 km^2^ composed by a mosaic of mature and secondary deciduous forest, surrounded by the Paranapanema River in the south, and by a matrix of cattle ranches and agriculture, mostly sugar cane, in the remaining borders (Uezu et al. 2008). Its average annual temperature is 22°C and annual rainfall is 1347 mm (Faria and Pires 2006). The park is part of the “Planalto Forest,” which is distinguished from the coastal forest of the Atlantic Forest biome by having lower annual rainfall and a marked dry season from May to September and is thus more similar to the Cerrado biome (Salis et al. 1995). In fact, the semi-deciduous forests of the “Planalto Forest” are similar to those occurring within or on the edges of the Cerrado (Salis et al. 1995).

#### Pantanal

Baía das Pedras Ranch, a private property of 145 km^2^, is located in the Nhecolândia Sub-Region of the Southern Pantanal, Municipality of Aquidauana (19°20’S, 55°43’W), Mato Grosso do Sul State, in the central-western region of Brazil. The ranch includes a mosaic of seasonally inundated grasslands, lakes, gallery forests, scrub, and deciduous forests that supports an abundance of wildlife and is situated far away from the edges of the biome where deforestation and other anthropogenic threats are occurring. Cattle are raised extensively on the native grasses. The average annual temperature is 25°C and annual rainfall is 1185 mm (Calheiros and Fonseca Júnior 1996).

#### Cerrado

The study site in the Cerrado biome is a 2200 km^2^ mosaic of private properties (cattle ranches and farms) and landless people settlements within the Municipalities of Nova Alvorada do Sul and Nova Andradina, Mato Grosso do Sul State (21°60’S, 53°83’W). The area includes small fragments of natural Cerrado habitat (Cerradão fragments, gallery forests, and marshland - 25% of the study area), surrounded by areas highly impacted by human activities such as agriculture (particularly sugarcane, soybean and corn), cattle-ranching (cultivated pastureland), Eucalyptus plantations, rural communities, and highways. The average annual temperature is 25°C and annual rainfall is 1185 mm.

In each study site, tapirs were captured by darting after physical restraint in either box traps or pitfall traps, or by darting from a distance (Quse et al. 2014). Animals were anesthetized mostly using a combination of butorphanol, medetomidine and ketamine, as described by Medici et al. (2014) and Fernandes-Santos et al. (2020). Reversal agents were administrated at the end of procedures. The procedures carried out during immobilization included the subcutaneous insertion of a microchip, morphometric measurements, sex and age class determination, physical examination, collection of biological samples for health and genetic studies, and placement of a telemetry collar on adults. Animals were tracked using VHF tracking (all three regions; Telonics® MOD500) and GPS tracking (Pantanal and Cerrado; Telonics® TGW SOB and GPS IRIDIUM models). A total of 74 tapirs were tracked starting in July of 1997 until October of 2019, with the majority of the individuals being in the Pantanal (46), while 17 and 11 were from the Cerrado and Atlantic Forest regions, respectively.

Tapirs equipped with VHF collars were monitored for 5 days per month with data collection concentrated during crepuscular times, 3 hours at dawn (04:00-07:00 h) and 3 hours at dusk (17:00-20:00 h). These periods are the two main peaks of tapir activity (Medici 2011). Each tapir was located every 30 minutes during the sampling periods. GPS collars were programmed to obtain a fix every hour and operated for a median of 15.4 months across all tagged tapirs. GPS fix success rates were 75% in the Pantanal and 90% in the Cerrado. The full dataset comprised 232,622 location estimates collected over a period of 22 years (for full details see Supplementary File S1). In addition to the tapir location data, we collected 883 and 174 measurements from tags in fixed locations in the Pantanal and Cerrado, respectively in order to calibrate the measurement error of the GPS tracking collars.

### Data analysis

Initial exploratory analyses were carried out in ctmmweb (version 0.2.11, Calabrese et al. 2021). All formal statistical analysis and plotting were performed using R (version 4.0.5, R Core Team 2021), with the packages ctmm (version 0.6.1, Calabrese et al. 2016), mgcv (version 1.8-36, Wood 2017), ggplot2 (version 3.3.4, Wickham 2016), ggmap (version 3.0.0, Kahle and Wickham 2013). The furrr package (version 0.2.2, Vaughan and Dancho 2021) was used for parallel computation on Windows machines. All R code can be found in the GitHub repository at https://github.com/StefanoMezzini/tapirs.

#### Data calibration and cleaning

Before analysis, we performed an error calibration and data cleaning process to minimise the impacts of GPS measurement error and outliers on our subsequent analyses (Fleming et al. 2020). Data cleaning and calibration were carried out using the methods implemented in the ctmm R package. For this process, location estimates collected via VHF telemetry were assumed to be free from any meaningful measurement error and raw locations were carried forward in the analyses. Measurement error on the GPS data was calibrated using a unitless Horizontal Dilution of Precision (HDOP), which quantifies the accuracy of each positional fix. We then estimated an equivalent range error with the HDOP values from the tags in fixed locations. This allowed for the unitless HDOP values to be converted into estimates of measurement error in meters. After calibration, data points were considered as outliers (and removed) if they had a large (error-informed) distance from the median location and the minimum speed required to explain the displacement was unusually high (≥ 1m/s). The Atlantic Forest dataset contained a total of 4,082 observations, 8 (ca. 0.2%) of which were removed as outliers; the Pantanal dataset contained 139,138 observations, 914 (ca. 0.7%) of which were removed; while the Cerrado dataset contained 90,402 observations, 193 (ca. 0.2%) of which were removed.

#### Movement modelling and home range estimation

For each of the monitored tapirs we quantified a number of key movement metrics and home range-related characteristics that allowed us to test for an effect of habitat composition and human disturbance on tapir movement behaviour. For this we first identified the best Continuous-Time Movement Model (CTMM) for each animal using the ctmm.select function from the ctmm package. This fits a series of CTMMs to location data using perturbative Hybrid Residual Maximum Likelihood (pHREML, Fleming et al. 2019) and chooses the best model using small-sample-sized corrected Akaike’s Information Criterion (AICc). The models used here are insensitive to sampling frequency (Johnson et al. 2008, Fleming et al. 2014, Blackwell et al. 2016) and they account for spatio-temporal autocorrelation in the data (when necessary), so they are robust to irregular or frequent sampling frequency (Fleming et al. 2018). The parameter estimates from each individual’s movement model provided information on the tapir’s home range crossing time (*τ* _*p*_, in days), and directional persistence timescale (*τ* _*V*_, in hours).

We then conditioned on the selected CTMMs to estimate each animal’s 95% home range (HR) area (in km) using small-sample-size bias corrected Autocorrelated Kernel Density Estimation (AKDE) (Fleming and Calabrese 2017), and average daily movement speed (in km/day) using continuous-time speed and distance (CTSD) estimation (Noonan et al. 2019).

#### Movement pattern analyses

We were first interested in understanding whether home-range areas and movement metrics differed across the three biomes, as well as between animals of different age and sex. For these comparisons, home range estimates were compared using the meta-analysis methods implemented in the ctmm package (Fleming et al. 2021), whereas other movement metrics were analysed using the meta-regression model implemented in the R package metafor (Viechtbauer 2010). This allowed for uncertainty in each individual estimate to be propagated into the population level estimate when making comparisons, minimising the risk of spurious significance.

To test whether tapirs responded to different environment types, the HR sizes and average daily speeds were regressed against the proportions of the habitat types in each HR. For the Atlantic Forest, we used the habitat map provided in the park’s management plan (Faria and Pires 2006). For the Pantanal and Cerrado, we obtained satellite imagery from the periods of data collection. Habitat classification was then carried out using GIS software, and a team of researchers confirmed the classifications in the field. Similarly, the HR sizes and average daily speeds were regressed against their HR’s average machine-learning-based human footprint index (ml-HFI) (Keys et al. 2021) to test whether human activity significantly altered the animals’ behavior. The ml-HFI is an index of human pressure on the landscape that is derived from remotely sensed surface imagery and ranges on a scale between 0 (no human impact), and 1 (high human impact). For these models we applied Generalized Additive Models (GAMs) with a Gamma distribution and a log link function for the response. The Gamma distribution allows for more accurate significance testing and is an appropriate distribution for variables that range between 0 and ∞, while the log link scale allows HFI to have a multiplicative effect on the response. The GAMs were fit using the mgcv package (Wood 2017) and Residual Maximum Likelihood (REML), and the best model was selected using AIC.

## Results

### Individual variation in movement and space use

The mean home range size across all monitored tapirs was 8.31 km (95% CI: 6.53 - 10.42; Fig. 2), ranging between 1 km^2^ and 29.7 km^2^ (Fig. 3a). Tapirs had HR crossing times of 0.72 days on average (95% CI: 0.35 - 1.10), ranging from 0.05 to 12.8 days (Fig. 3b), and a mean velocity autocorrelation timescale of 0.44 hours (95% CI: 0.39 - 0.49), ranging from 0.17 to 1.88 hours (Fig. 3c). We estimated that tapirs had mean movement speeds of 11.2 km/day (95% CI: 10.2 - 12.1), ranging from 1.51 to 25.96 km/day (Fig. 3d). There was no evidence that average daily speed differed between sexes (females: 10.5 km/day, 95% CI: 9.19 - 12.0; males: 11.9 km/day; 95% CI: 10.3 - 13.7, *p*= 0.22, 4a), nor between age groups (adults: 11.8 km/day, 95% CI: 10.6 - 13.2; sub-adults: 9.52 km/day, 95% CI: 7.94 - 11.4;, Fig. 4b).

**Figure 2:**
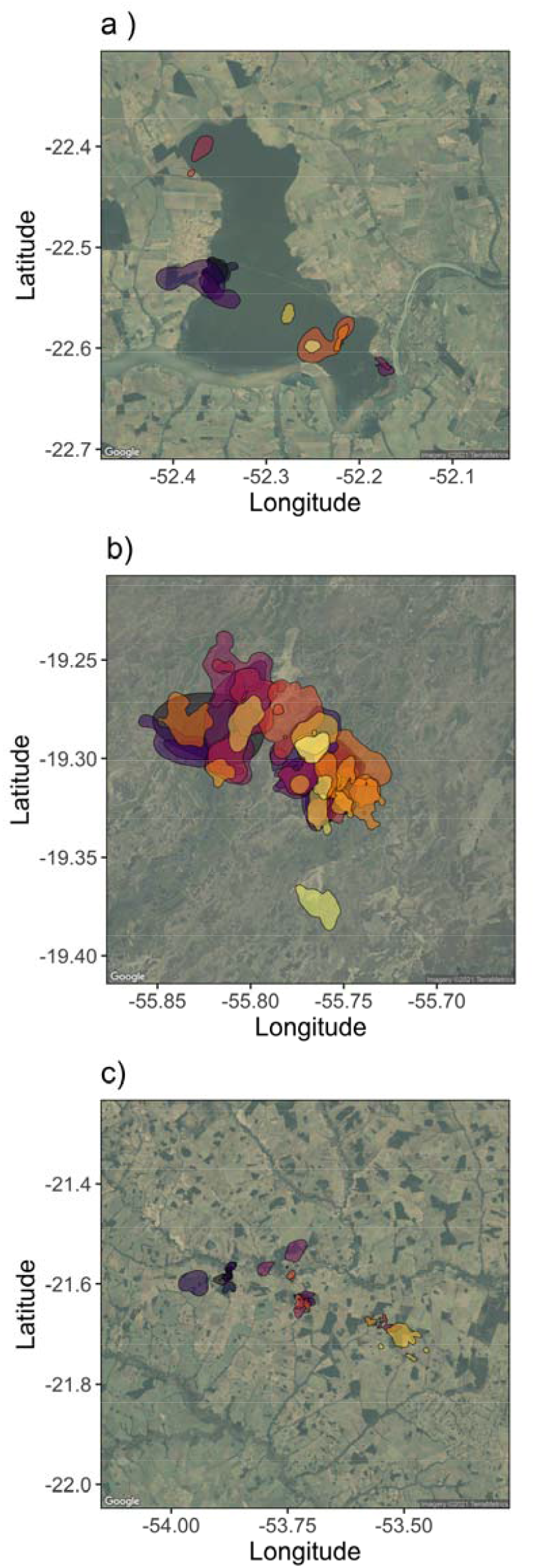
Autocorrelated kernel density estimates of each tapir’s 95% home range in each of the three regions: a) Atlantic Forest, b) Pantanal, and c) Cerrado.

**Figure 3:**
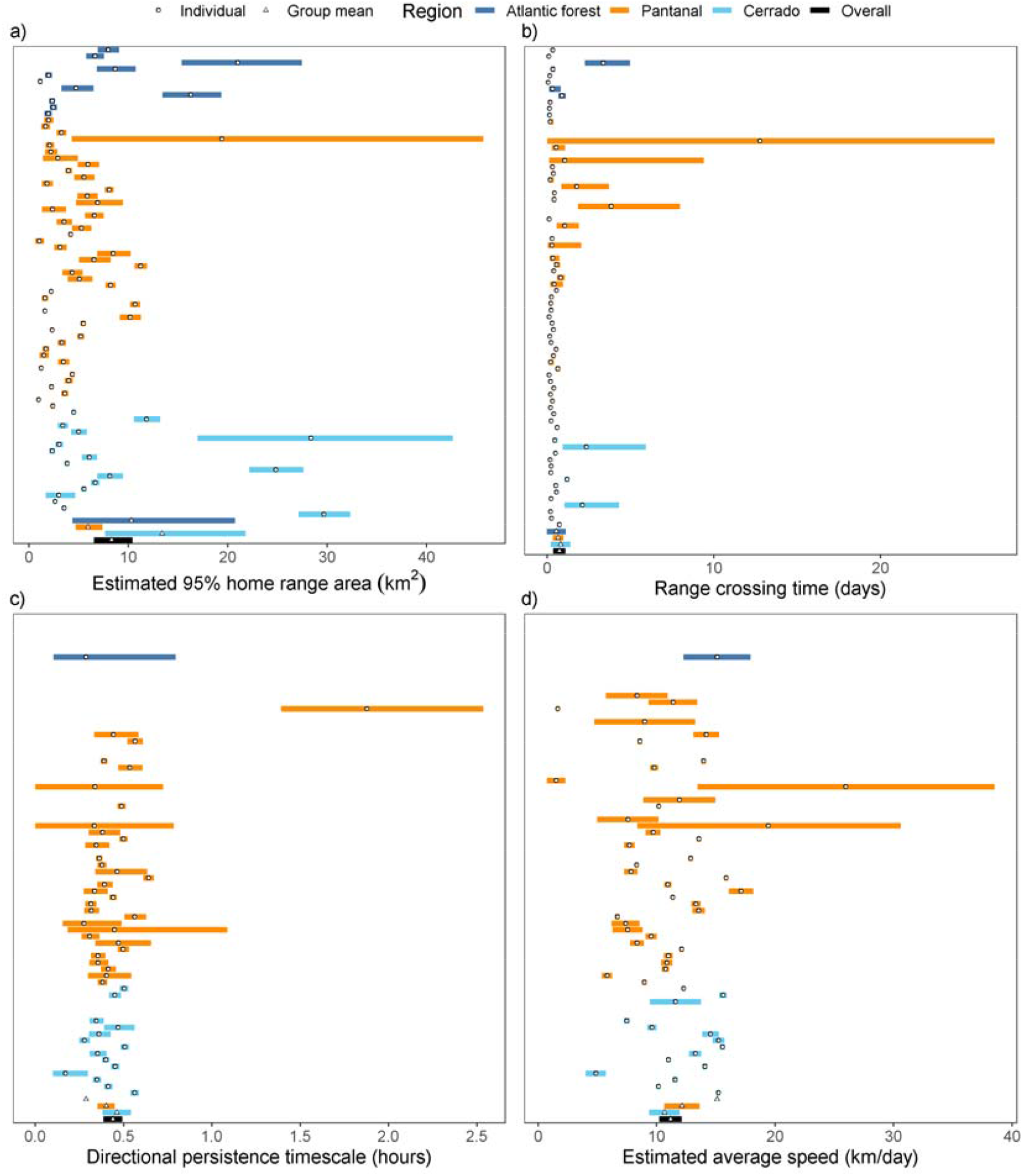
Parameter estimates from each tapir’s movement model (circles) and group means (triangles), with 95% confidence intervals. Individuals with a movement model that does not allow for inferences in movement speed are left blank.

**Figure 4:**
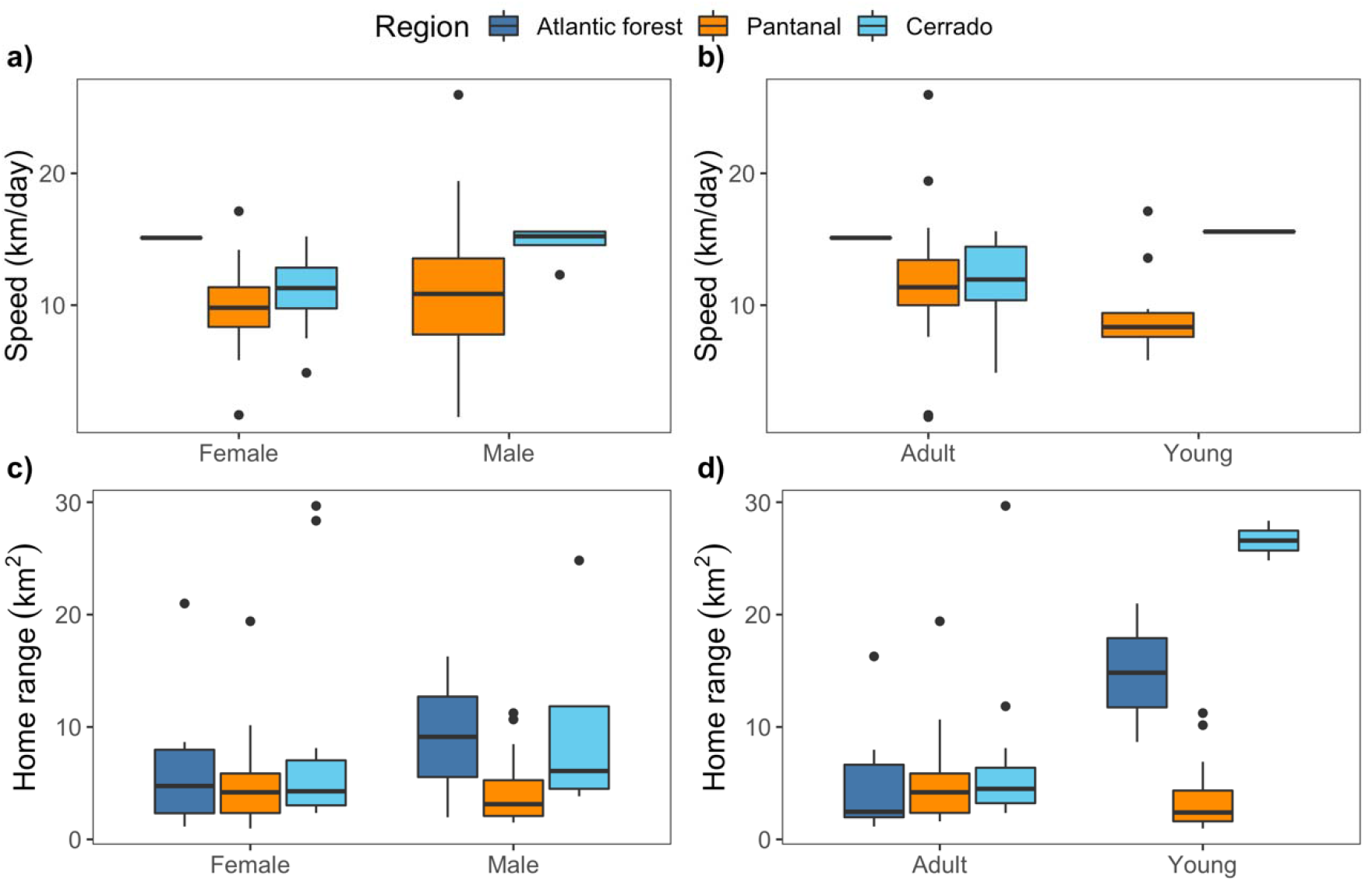
Boxplots of daily average speed (a, b) and estimated home range size (c, d) by sex and age group. We note that estimation of movement speeds for adult females was only possible for a single tapir in the Atlantic Forest. In addition, we could only estimate speed for a single young tapir in the Cerrado.

There was no evidence that home range sizes differed between sexes (males: 5.43 km^2^, 95% CI: 3.84 - 7.68; females: 6.27 km^2^, 95% CI: 4.64 - 8.48; p= 0.541, Fig. 4c) nor between age groups (adults: 5.47 km^2^, 95% CI: 4.21 - 7.1; sub-adults: 7.01 km^2^, 95% CI: 4.63 - 10.6; p = 0.324, Fig. 4d).

### Variation in movement across biomes and gradients of human disturbance

The Atlantic Forest, Cerrado, and Pantanal vary substantially in habitat composition, levels of human disturbance, and tapir population densities. Despite these differences, we found that lowland tapir movement behaviour and space use were consistent across all three biomes (Fig. 3).

We also found that habitat type had little effect on HR area or average individual movement speeds. The best HR area regression model only accounted for the effect of areas of exposed soil (approximate p-value: 0.023, = 0.477; Fig. 5a), while no land use types had a significant effect on an animal’s average speed. There was very little difference between the AIC of the full model (315.69, df = 10.18, 7 predictors and an intercept) and that of the intercept-only model (310.89, df = 2). However, the directional persistence term () was significantly lower for animals who had a higher amount of forested area (Fig. 5b) or water (Fig. 5c) in their home ranges. Importantly, we note here that the significant differences in directional persistence persisted even after adjusted for the increased location error in the forested areas.

**Figure 5:**
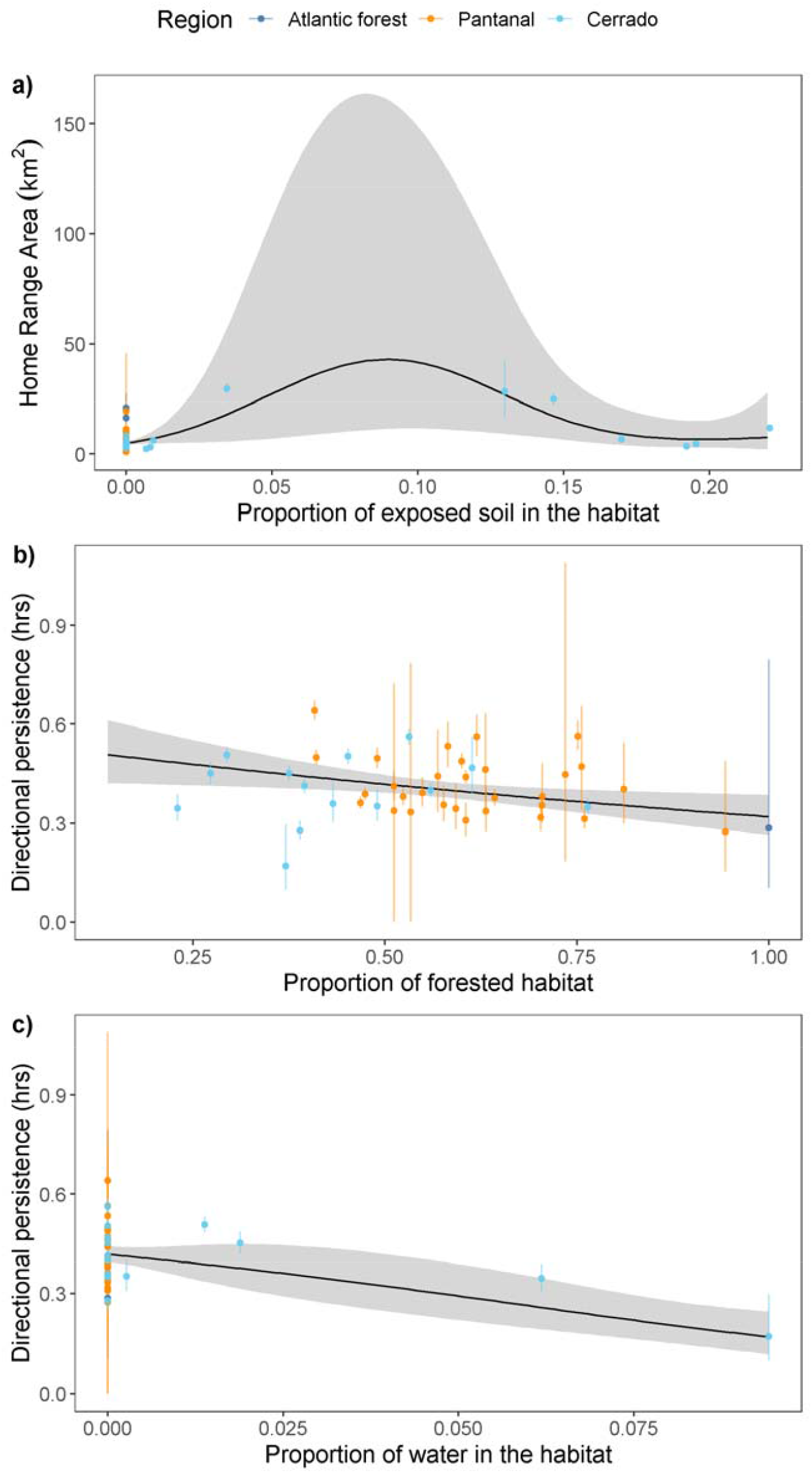
Effect of habitat types on lowland tapir space use and movement. The vertical error-bars indicate the 95% confidence intervals for the movement parameter estimates. Panel a) depicts the estimated mean effect of exposed soil on the tapirs’ estimated home range area. The effects of b) forested area or c) water in a tapir’s home range on its estimated directional persistence are also shown.

HFI had no significant effect on lowland tapir home range size (p-value = 0.90; Fig. 6a), nor average daily movement speed (p-value = 0.53; Fig. 6b), nor directional persistence (p-value = 0.596, 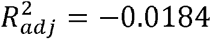). A tapir living in a near pristine environment (HFI = 0.004) had a home range estimate of 7.77 km (95% CI: 2.12 - 28.6) and an average speed of 13.19 km/day (95% CI: 7.82 - 22.1) with a directional persistence of 0.355 hours (95% CI: 0.160 - 0.784), while a tapir from the most altered habitat we monitored (HFI = 0.31) had an estimated home range area of 6.93 km (95% CI: 3.36 - 14.3) and an average speed of 10.43 km/day (95% CI: 8.27 - 13.2) with a directional persistence of 0.478 hours (95% CI: 0.335 - 0.683).

**Figure 6:**
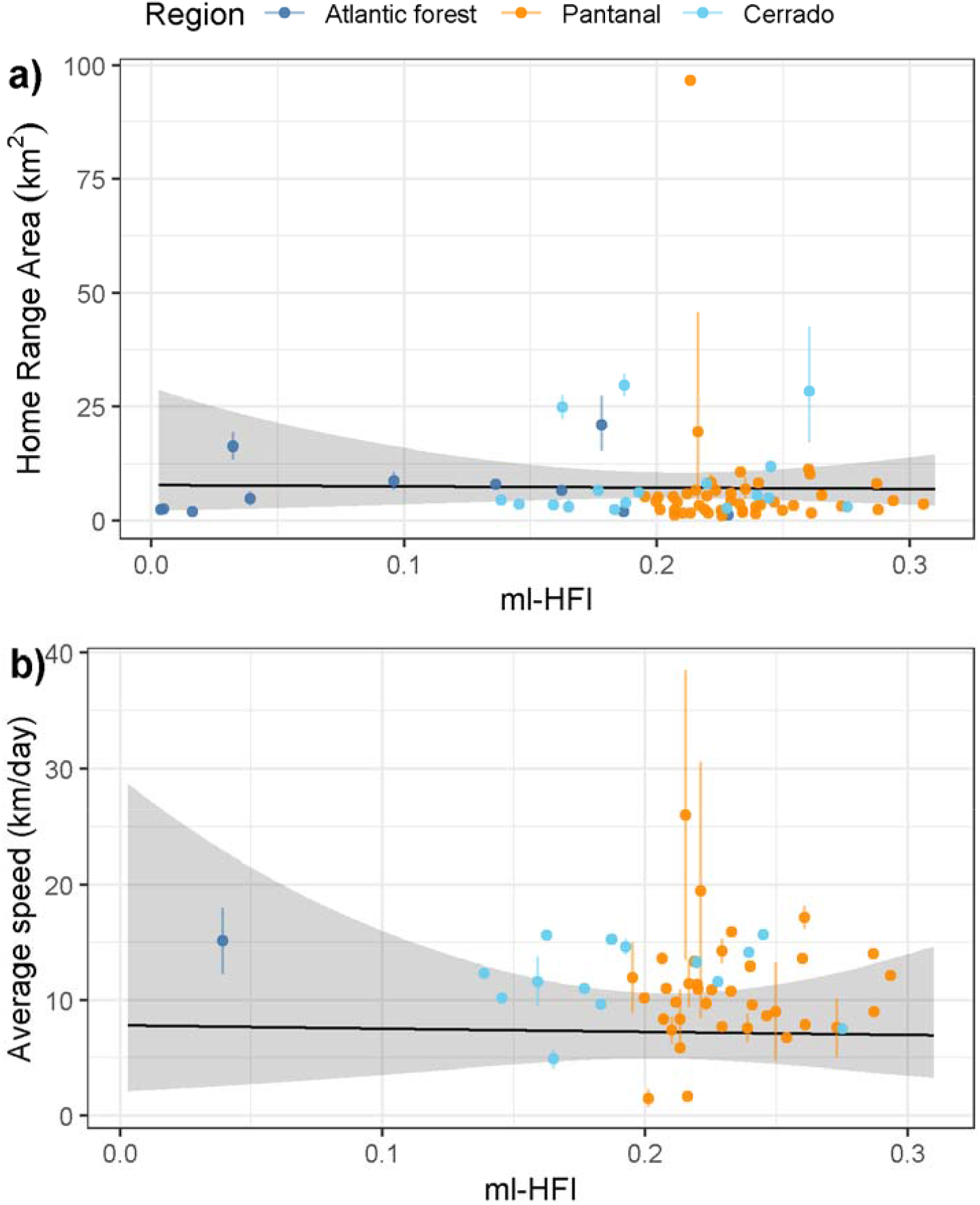
Estimated mean effect of machine-learning-based human footprint index (ml-HFI) on the tapirs’ estimated home range area and estimated average daily speed. The vertical segments indicate the 95% confidence intervals for the movement parameter estimates.

## Discussion

Understanding individual movement and space use requirements is a key step in conservation planning (Allen and Singh 2016). Prior to the present study, very little was known about the movement ecology of tapirs despite their vulnerable status and declining population sizes (Varela et al. 2019). From detailed tracking of 74 tapirs collected over 22 years, we found that although individuals varied in their movement, these inter-individual differences were not explained by differences in age, sex, habitat composition, biome, nor human disturbance. Overall, human activity and land use change did not appear to significantly affect their behaviour one way or another. This contradicts patterns in large herbivores generally (Tucker et al. 2018; Doherty et al. 2021), and further emphasizes the need to understand the movement ecology of target populations when designing conservation and recovery strategies.

### The ecology of lowland tapir space use

Interestingly, we found that the home range sizes and mean daily movement speeds of lowland tapirs were consistent across the three study sites. This consistency in movement was surprising as these different biomes have substantially different habitat compositions, patterns of seasonality, and productivity (Morato et al. 2016; see also Appendix S1). Tapirs living in the Pantanal, for instance, occupy a near pristine ecosystem but must cope with significant seasonal flooding, whereas individuals in the Cerrado occupied an agricultural and cattle ranching mosaic with more stability across seasons. The unique requirements of these three different biomes, however, did not impact the space use and movement speed of tapirs in any statistically detectable way. Furthermore, the only pre-existing study on tapir movement found that individuals had complex home range structures, with multiple core areas of use that were established according to the distribution of patches of preferred habitat types (Tobler 2008). While individuals may exhibit differential use of patchily distributed resources, we found that habitat composition had no effect on home range sizes. In addition to exhibiting little inter-individual variation in movement, variogram analysis (Fleming et al. 2014) showed that tapir movement was extremely consistent over time (see also Fleming et al. 2019). Here again, this seasonal stability in movement was interesting, especially for animals living in the Pantanal where, every year, large parts of the biome change from terrestrial into aquatic habitats and vice-versa (Alho 2008). We note though that the flooding regime in the Pantanal has been changing over the last decade and the biome is expected to become drier under the IPCC’s climate change scenarios (Marengo et al. 2015).

We did find that animals with a higher proportion of forest and/or more water bodies in their home ranges had reduced directional persistence. This shows how habitat complexity can impact movement (Dickie et al. 2017), with potential implications for foraging efficiency and encounter rates (Visser and Kiørboe 2006; Bartumeus et al. 2008; Martinez-Garcia et al. 2020). Nonetheless, these differences did not translate into patterns in tapir home range sizes and mean daily movement speeds.

### Lowland tapir movement across a gradient of human disturbance

This is the first study aimed at understanding how lowland tapir space use and movement vary across differing biomes and degrees of human disturbance. Contrary to our initial expectations, and to patterns in large herbivores generally (Doherty et al. 2021), human impacts on the landscape also had no measurable effect on tapir movement. Tapirs inhabiting the Atlantic Forest, the most disturbed biome with only 12.4% of habitat remaining (SOS Mata Atlântica 2008), had home range sizes that were comparable in size to tapirs inhabiting the Cerrado, a biome that has lost almost 50% of its natural area (Machado et al. 2004; Ministério da Agricultura 2021), and the Pantanal, a near pristine biome. Notably, the Lowland Tapir Conservation Action Plan published by the IUCN SSC Tapir Specialist Group (TSG) in 2007 (Medici et al. 2007), and the Lowland Tapir National Action Plan (PAN – Plano de Ação Nacional, ICMBIO – Instituto Chico Mendes de Conservação da Biodiversidade, Brazil) published in 2019 prioritize the mitigation of the impacts of small, isolated tapir populations. Population isolation thus emerges as one of the most important threats to the species’ long-term persistence. However, addressing this issue will require additional efforts as the average and maximum distances we recorded for tapir movements were substantially less than the distances between most tapir populations.

Humans are directly responsible for more than one-quarter of global terrestrial vertebrate mortality (Hill et al. 2019). Mortality at this scale is expected to impose strong selection pressure on animal populations (Oro et al. 2013; Swaddle et al. 2015). As genotypic adaptation takes generations to occur (Barnosky and Kraatz 2007), behavioral plasticity provides the most immediate response to the pressures of Human Induced Rapid Environmental Change (HIREC, Sih et al. 2011). The capacity for behavioural plasticity in movement and space use in response to human disturbance is especially important for long-lived, K-selected species such as tapirs (Rosenheim and Tabashnik 1991; Sih et al. 2011; Montgomery et al. 2020) that take years to reach sexual maturity and have long inter-generational intervals (Medici 2011). Despite the key importance of behavioural adaptations in response to HIREC, tapir movement appeared to exhibit very little plasticity in response to human disturbance. The lack of any measurable response to human activity suggests that tapirs living near humans may experience increased exposure to vehicle collisions (Medici 2019; Abra et al. 2020), pesticide and environmental pollutants (Medici et al. 2014; Fernandes-Santos et al. 2020; Medici et al. 2021) and poaching (Sanches et al. 2011). Human modified habitats thus risk being ecological traps (Schlaepfer et al. 2002) for tapirs as individuals showed no detectable responses to degradations in habitat quality. Although tapir home range area and mean daily movement speed exhibited no statistically detectable response to the human footprint index, it is possible that individuals are responding to human disturbance at a finer temporal and/or spatial scale than the long-term averages that were examined here. It may also be possible that tapirs exhibit non-linear, or even binary, responses to human disturbance that were not possible to detect. Future investigation into lowland tapir behaviour in more heavily modified habitats is clearly warranted.

## Conclusions

We compared home range areas and movement behavior of lowland tapirs using telemetry data collected over 22 years across 3 biomes in southern Brazil: the Pantanal, Cerrado, and Atlantic Forest. These data represent the largest lowland tapir tracking dataset yet to be collected, with over 232,000 locations from 74 tracked individuals and fill a critical knowledge gap in lowland tapir ecology, which can contribute to long-term species management and conservation planning. Contrary to our expectations, we observed very little individual variability in lowland tapir space use and movement, and human impacts on the landscape also had no measurable effect on their movement. Lowland tapir movement behaviour thus appears to exhibit very little phenotypic plasticity. The lack of any adaptive response to anthropogenic disturbance suggests that human modified habitats risk being ecological traps for tapirs and this information should be factored into conservation actions aimed towards protecting lowland tapir populations.

## Acknowledgments

The study of tapir movement ecology has been an important component of the long-term activities of the Lowland Tapir Conservation Initiative (LTCI) – Instituto de Pesquisas Ecológicas (IPÊ) in Brazil. The LTCI has the institutional support from the International Union for Conservation of Nature (IUCN) Species Survival Commission (SSC) Tapir Specialist Group (TSG), Association of Zoos and Aquariums (AZA) Tapir Taxon Advisory Group (TAG), and European Association of Zoos and Aquariums (EAZA) Tapir Taxon Advisory Group (TAG). LTCI’s financial support comes from national and international agencies, including zoological institutions, foundations, private businesses, and private individuals. MJN was supported by an NSERC Discovery Grant RGPIN-2021-02758. This work was partially funded by the Center of Advanced Systems Understanding (CASUS) which is financed by Germany’s Federal Ministry of Education and Research (BMBF) and by the Saxon Ministry for Science, Culture and Tourism (SMWK) with tax funds on the basis of the budget approved by the Saxon State Parliament. CHF and JMC were supported by NSF IIBR 1915347. EPM would like to thank the Smithsonian Conservation Biology Institute (SCBI) for hosting her for a 2-month research visit for initial data processing and analysis.

## Appendix S1

In this appendix we provide supporting information on all of the analyses presented in the main text.

### Home range estimates and data collection methods

The tapir location data included in this study were collected over 22 years using a mix of VHF and GPS tracking. As these two data types resulted in different sampling frequencies, it was possible for differences in autocorrelation to drive differences in the estimated home range areas (Noonan et al. 2019). We therefore carried out a supporting analysis to ensure that there was no relationship between data collection methods and the estimated home range estimates. We found that there was no significant difference in the home range estimates between individuals who were monitored using GPS collars, VHF tracking, or a mixture of the two (GPS as the control, p-values: 0.495 for GPS and VHF, 0.739 for VHF only, see Fig. S1).

**Figure S1:**
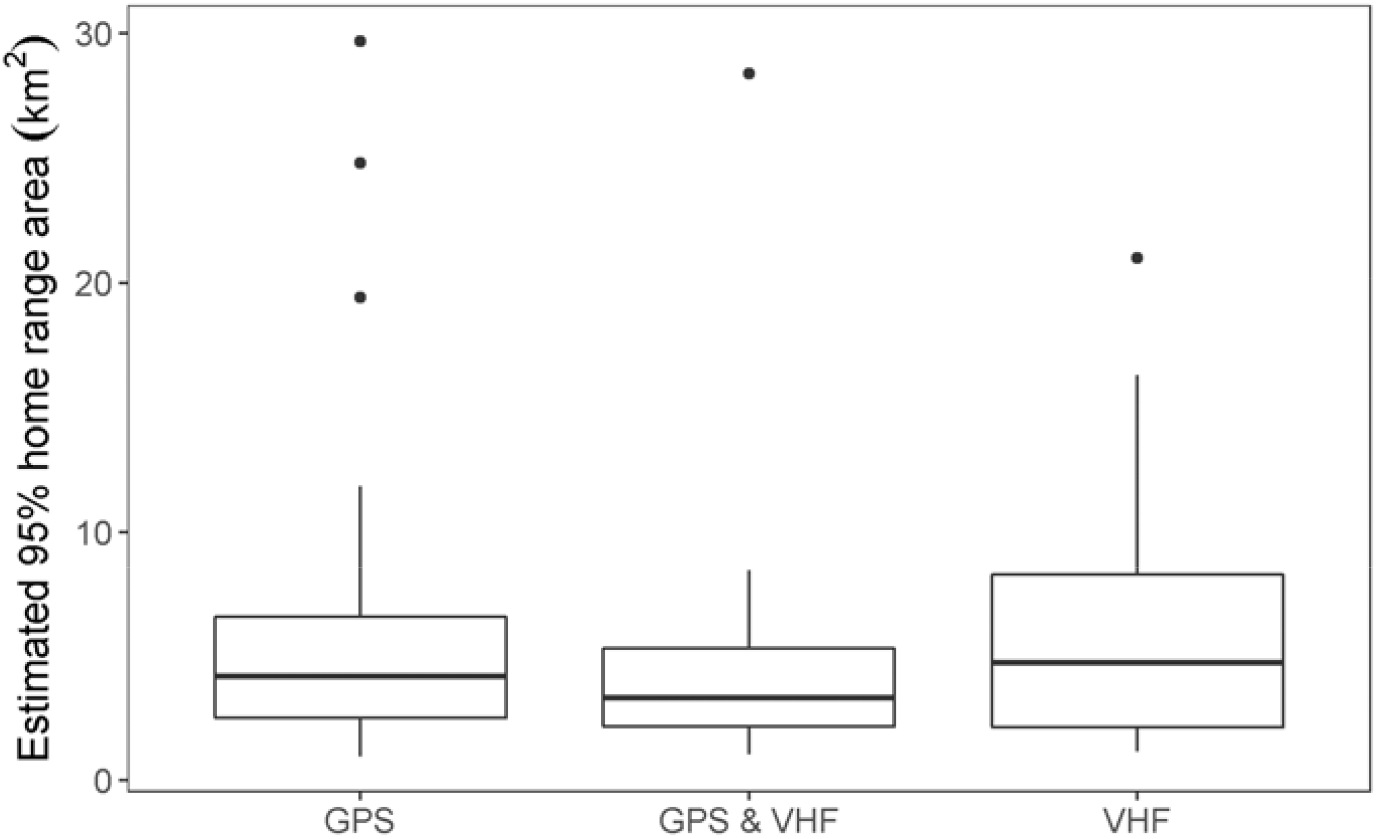
Estimated 95% home range area by tracking method. Tapirs were monitored using GPS collars, Very High Frequency (VHF) tracking, or both.

These findings suggest that any of the results presented in the main text are robust to inter-individual differences in data collection.

### Habitat composition

Here we show how the habitat composition differed between each of the three study areas. In addition, we show how the proportion of each land use type within the home range of each tapir.

**Figure S2:**
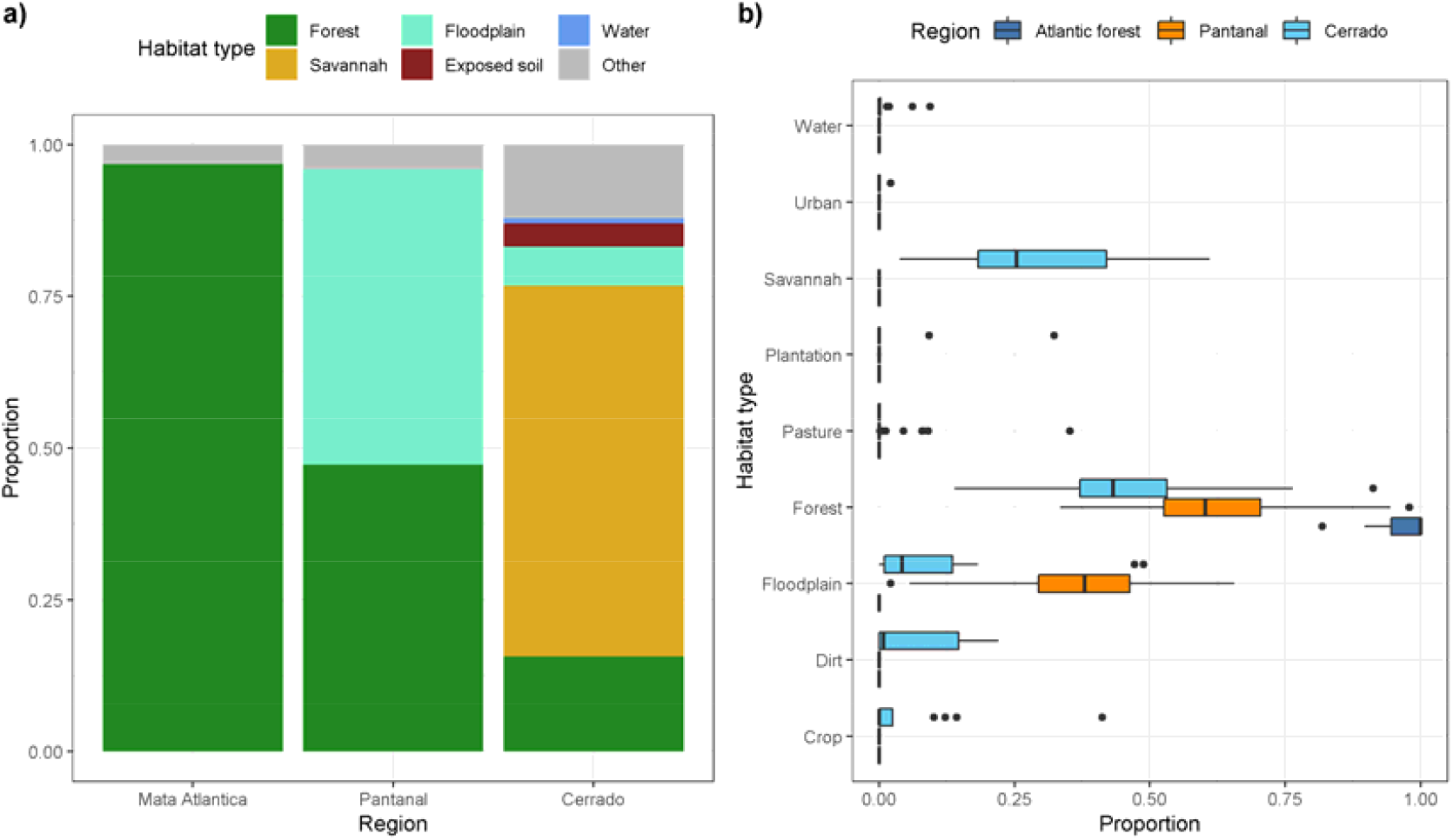
Figure depicting a) the proportion of habitat type in each of the three study areas, and b) the proportion of each land use type within the home range of each tapir.

### Influence of outliers

As noted in the main text, HFI had no significant effect on lowland tapir home range size, nor average daily movement speed, nor directional persistence. These findings were consistent before and after removing outliers (p-value = 0.596,; p-value = 0.188,, respectively, Fig. S3).

**Figure S3:**
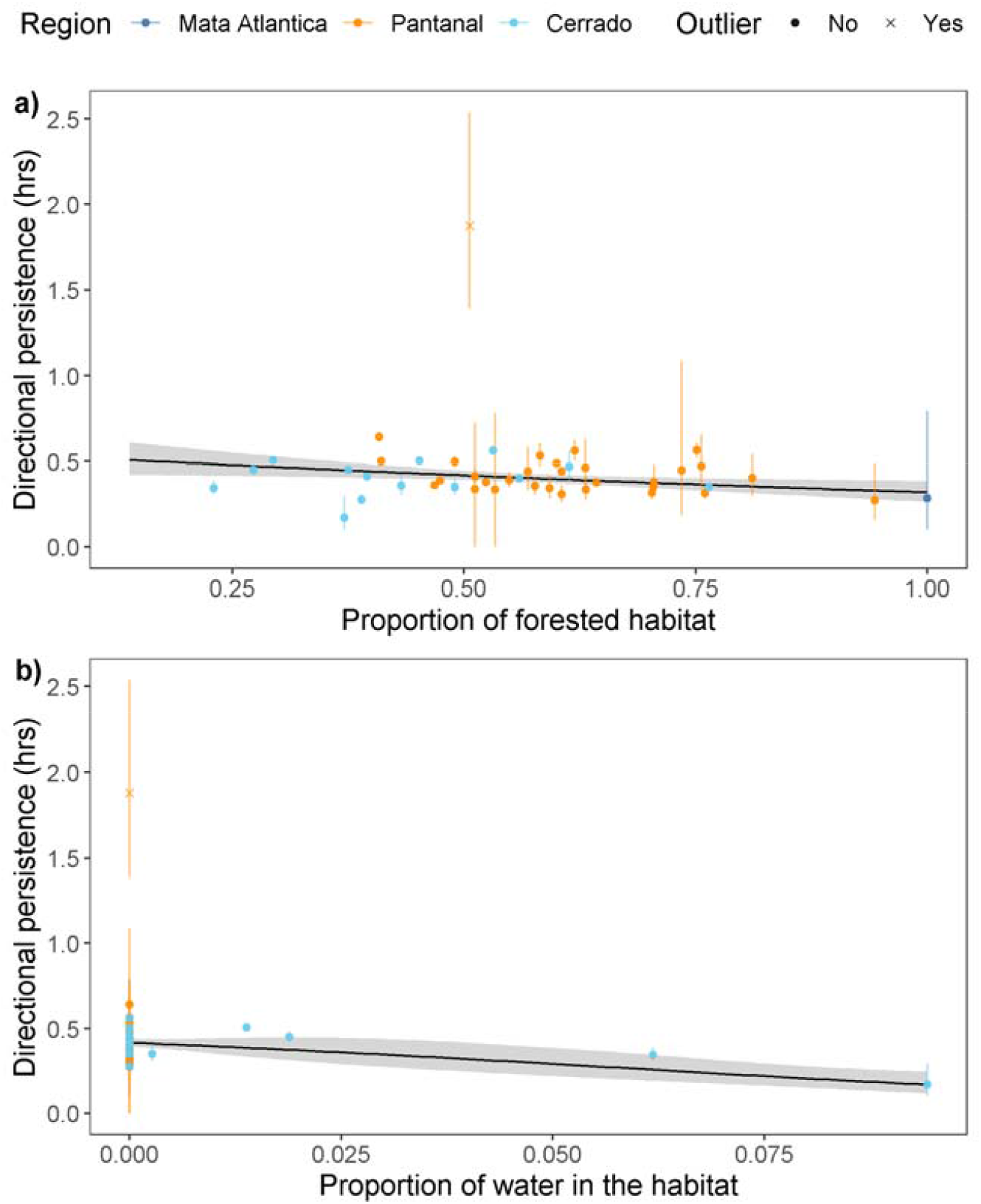
Effect of forested area or water in a tapir’s home range on its estimated directional persistence. The point indicated as an outlier was removed from the dataset for the purpose of the regression.

## Notes

https://github.com/StefanoMezzini/tapirs

### Competing Interest Statement

The authors have declared no competing interest.

## References

Alho CJR. Biodiversity of the Pantanal: Response to seasonal flooding regime and to environmental degradation. Brazilian Journal of Biology. 2008; DOI:10.1590/S1519-69842008000500005

Allen AM, Navinder JS. Linking movement ecology with wildlife management and conservation. Frontiers in Ecology and Evolution. 2016; DOI:10.3389/fevo.2015.00155

Barnosky, AD, Kraatz BP. The role of climatic change in the evolution of mammals. BioScience. 2007; DOI:10.1641/B570615

Bartumeus F, Catalan J, Viswanathan GM, Raposo EP, Da Luz MGE. The influence of turning angles on the success of non-oriented animal searches. Journal of Theoretical Biology. 2008; DOI:10.1016/j.jtbi.2008.01.009

Bauer S, Hoye BJ. Migratory animals couple biodiversity and ecosystem functioning worldwide. Science. 2014; DOI:10.1126/science.1242552

Blackwell PG, Niu M, Lambert MS, LaPoint SD. Exact Bayesian inference for animal movement in continuous time. Methods in Ecology and Evolution. 2016; DOI:10.1111/2041-210X.12460

Brooks TM, Mittermeier RA, Mittermeier CG, Fonseca GAB, Rylands AB, Konstant WR, et al. Habitat loss and extinction in the hotspots of biodiversity. Conservation Biology. 2002; DOI:10.1046/j.1523-1739.2002.00530.x

Calabrese JM, Fleming CH, Gurarie E. Ctmm: An r Package for analyzing animal relocation data as a continuous-time stochastic process. Methods in Ecology and Evolution. 2016; DOI:10.1111/2041-210X.12559

Calabrese JM, Fleming CH, Noonan MJ, Dong X. Ctmmweb: a graphical user interface for autocorrelation-informed home range estimation. Wildlife Society Bulletin. 2021; DOI:10.1002/wsb.1154

Calder WA III. Ecological scaling: mammals and birds. Annual Review of Ecology and Systematics. 1983;14:213–230.

Calheiros DF, Fonseca Junior WC. Perspectivas de estudos ecologicos sobre o Pantanal. Empresa Brasileira de Pesquisa Agropecuaria (EMBRAPA-CPAP). Corumba, Mato Grosso do Sul, Brazil. 1996.

Cardinale BJ, Duffy JE, Gonzalez A, Hooper DU, Perrings C, Venail P, et al. Biodiversity loss and its impact on humanity. Nature. 2012; DOI:10.1038/nature11148

Cosgrove AJ, McWhorter TJ, Maron M. Consequences of impediments to animal movements at different scales: a conceptual framework and review. Diversity and Distributions. 2018; DOI:10.1111/ddi.12699

Dickie M, Serrouya R, McNay RS, Boutin S. Faster and farther: wolf movement on linear features and implications for hunting behaviour. Journal of Applied Ecology. 2017; DOI:10.1111/1365-2664.12732

Doherty TS, Hays GC, Driscoll DA. Human disturbance causes widespread disruption of animal movement. Nature, Ecology & Evolution. 2021; DOI:10.1038/s41559-020-01380-1

Esfahani HS, Ramírez MT. Institutions, infrastructure, and economic growth. Journal of Development Economics. 2003; DOI:10.1016/S0304-3878(02)00105-0

Fahrig L. Relative effects of habitat loss and fragmentation on population extinction. The Journal of Wildlife Management. 1997;61(3):603–610.

Fahrig L. Non-optimal animal movement in human-altered landscapes. Functional Ecology. 2007; DOI:10.1111/j.1365-2435.2007.01326.x

Fernandes-Santos RC, Medici EP, Testa-Jose C, Micheletti T. Health assessment of wild lowland tapirs (Tapirus terrestris) in the highly threatened Cerrado biome, Brazil. Journal of Wildlife Diseases. 2020; DOI:10.7589/2018-10-244

Fleming CH, Drescher-Lehman J, Noonan MJ, Akre TSB, Brown DJ, Cochrane MM, et al. A comprehensive framework for handling location error in animal tracking data. Ecology. 2020. Preprint.

Fleming CH, Noonan MJ, Medici EP, Calabrese JM. Overcoming the challenge of small effective sample sizes in home-range estimation. Methods in Ecology and Evolution. 2019; DOI:10.1111/2041-210X.13270

Fleming CH, Sheldon D, Fagan WF, Leimgruber P, Mueller T, Nandintsetseg D, et al. Correcting for missing and irregular data in home-range estimation. Ecological Applications. 2018; DOI:10.1002/eap.1704

Fleming CH, Calabrese JM, Mueller T, Olson KA, Leimgruber P, Fagan WF. From fine-scale foraging to home ranges: a semivariance approach to identifying movement modes across spatiotemporal scales. The American Naturalist. 2014; DOI:10.1086/675504

Fleming CH, Calabrese JM. A new kernel density estimator for accurate home-range and species-range area estimation. Methods in Ecology and Evolution. 2017; DOI:10.1111/2041-210X.12673

Fleming CH, Deznabi I, Alavi S, Crofoot MC, Hirsch BT, Medici EP, et al. Population-level inference for home-range areas. bioRxiv – The Preprint Server for Biology. 2021; DOI:10.1101/2021.07.05.451204

Gardner AL. Mammals of South America, Volume 1: Marsupials, Xenarthrans, Shrews, and Bats. University of Chicago Press; 2008.

Peng H, Montiglio PO, Somveille M, Cantor M, Farine DR. The role of habitat configuration in shaping animal population processes: a framework to generate quantitative predictions. Oecologia. 2021; DOI:10.1007/s00442-021-04967-y

Faria H, Pires AS. Parque Estadual Morro Do Diabo - Plano de Manejo. Governo do Estado de Sao Paulo, Secretaria do Meio Ambiente, Instituto Florestal. Santa Cruz do Rio Pardo, Sao Paulo, Brazil. 2006.

Hill JE, DeVault TL, Belant JL. Cause-specific mortality of the world’s terrestrial vertebrates. Global Ecology and Biogeography. 2019; DOI:10.1111/geb.12881

Hill JE, DeVault TL, Wang G, Belant JL. Anthropogenic mortality in mammals increases with the human footprint. Frontiers in Ecology and the Environment. 2020; DOI:10.1002/fee.2127

Hooper DU, Adair EC, Cardinale BJ, Byrnes JEK, Hungate BA, Matulich KL, et al. A global synthesis reveals biodiversity loss as a major driver of ecosystem change. Nature. 2012; DOI:10.1038/nature11118

Kahle D, Wickham H. Ggmap: spatial visualization with Ggplot2. The R Journal. 2013;5(1):144.

Keys PW, Barnes EA, Carter NH. A machine-learning approach to human footprint index estimation with applications to sustainable development. Environmental Research Letters. 2021; DOI:10.1088/1748-9326/abe00a

Lucherini M, Lovari S. Habitat richness affects home range size in the red fox Vulpes vulpes. Behavioural Processes. 1996; DOI:10.1016/0376-6357(95)00018-6

Machado RB, Ramos-Neto MB, Pereira PGP, Caldas EF, Goncalves DA, Santos NS, et al. Estimativas de perda da area do cerrado brasileiro. Relatorio Tecnico. Conservacao Internacional, Brasilia, DF, Brazil. 2004.

Marengo JA, Oliveira GS, Alves LM. Climate change scenarios in the Pantanal. In: Bergier I, Assine ML, editors. Dynamics of the Pantanal Wetland in South America. Springer; 2016. p 227–238.

Martinez-Garcia R, Fleming CH, Seppelt R, Fagan WF, Calabrese JM. How range residency and long-range perception change encounter rates. Journal of Theoretical Biology. 2020; DOI:10.1016/j.jtbi.2020.110267

Medici EP. Family Tapiridae (Tapirs). In: Wilson DE, Mittermeier RA, editors. Handbook of the Mammals of the World: Volume 2: Hoofed Mammals. Lynx Edicions Barcelona; 2011. p 182–204.

Medici EP, Mangini PR, Fernandes-Santos RC. Health assessment of wild lowland tapir (Tapirus terrestris) populations in the Atlantic Forest and Pantanal Biomes, Brazil (1996–2012). Journal of Wildlife Diseases. 2014; DOI:10.7589/2014-02-029

Medici EP, Abra FD. Licoes aprendidas na conservacao da anta brasileira e os desafios para mitigar uma de suas ameacas mais graves: o atropelamento em rodovias. Boletim da Sociedade Brasileira de Mastozoologia. 2019;85:152–160.

Medici EP, Desbiez ALJ, Goncalves da Silva A, Jerusalinsky L, Chassot O, Montenegro OL, et al. Lowland Tapir (Tapirus terrestris) Conservation Workshop: Final Report. IUCN SSC Tapir Specialist Group & IUCN SSC Conservation Planning Specialist Group (Brazil Network). 2007.

Medici EP, Fernandes-Santos RC, Testa-Jose C, Godinho AF, Brand AF. Lowland tapir exposure to pesticides and metals in the Brazilian Cerrado. Wildlife Research. 2021; DOI:10.1071/WR19183

Ministerio da Agricultura, Pecuaria e Abastecimento. TerraClass - Gestao Integrada da Paisagem no Bioma Cerrado. 2021.

Montgomery RA, Macdonald DW, Hayward MW. The inducible defences of large mammals to human lethality. Functional Ecology. 2020; DOI:10.1111/1365-2435.13685

Morato RG, Stabach JA, Fleming CH, Calabrese JM, Paula RC, Ferraz KMPM, et al. Space use and movement of a neotropical top predator: the endangered jaguar. PLoS ONE. 2016; DOI:10.1371/journal.pone.0168176

Nilsen EB, Herfindal I, Linnell JDC. Can intra-specific variation in carnivore home-range size be explained using remote-sensing estimates of environmental productivity? Ecoscience. 2005; DOI:10.2980/i1195-6860-12-1-68.1

Noonan MJ, Tucker MA, Fleming CH, Alberts SC, Ali AH, Altmann J, et al. A comprehensive analysis of autocorrelation and bias in home range estimation. Ecological Monographs. 2019; DOI:10.1002/ecm.1344

Noonan MJ, Fleming CH, Akre TS, Drescher-Lehman J, Gurarie E, Harrison AL, et al. Scale-insensitive estimation of speed and distance traveled from animal tracking data. Movement Ecology. 2019; DOI: 1186/s40462-019-0177-1

Noonan MJ, Fleming CH, Tucker MA, Kays R, Harrison AL, Crofoot MC, et al. Effects of body size on estimation of mammalian area requirements. Conservation Biology. 2020; DOI:10.1111/cobi.13495

Noss AJ, Cuellar RL, Barrientos J, Maffei L, Cuellar E, Arispe R, et al. A camera trapping and radio telemetry study of lowland tapir (Tapirus terrestris) in Bolivian dry forests. Plant Diversity. 2003;229:44–45.

Oro D, Genovart M, Tavecchia G, Fowler MS, Martinez-Abrain A. Ecological and evolutionary implications of food subsidies from humans. Ecology Letters. 2013; DOI:10.1111/ele.12187

Powers RP, Jetz W. Global habitat loss and extinction risk of terrestrial vertebrates under future land-use-change scenarios. Nature Climate Change. 2019; DOI:10.1038/s41558-019-0406-z

Quse VB, Fernandes-Santos RC. Tapir Veterinary Manual. IUCN SSC Tapir Specialist Group (TSG). 2014.

R Core Team. R: a language and environment for statistical computing. Vienna, Austria: R Foundation for Statistical Computing. https://www.R-project.org. 2021.

Relyea RA, Lawrence RK, Demarais S. Home range of desert mule deer: testing the body-size and habitat-productivity hypotheses. The Journal of Wildlife Management. 2000;64(1):146–153.

Rosenheim JA, Tabashnik BE. Influence of generation time on the rate of response to selection. The American Naturalist. 1991;137(4):527–541.

Salis SM, Shepherd GJ, Joly CA. Floristic comparison of mesophytic semideciduous forests of the interior of the state of Sao Paulo, Southeast Brazil. Vegetatio. 1995;119(2):155–164.

Sanches A, Perez WAM, Figueiredo MG, Rossini BC, Cervini M, Galetti PM, et al. Wildlife forensic DNA and lowland tapir (Tapirus terrestris) poaching. Conservation Genetics Resources. 2011; DOI:10.1007/s12686-010-9318-y

Schick RS, Loarie SR, Colchero F, Best BD, Boustany A, Conde DA, et al. Understanding movement data and movement processes: current and emerging directions. Ecology Letters. 2008; DOI: 10.1111/j.1461-0248.2008.01249.x

Schlaepfer MA, Runge MC, Sherman PW. Ecological and evolutionary traps. Trends in Ecology & Evolution. 2002; DOI:10.1016/S0169-5347(02)02580-6

Sih A, Ferrari MCO, Harris DJ. Evolution and behavioural responses to human-induced rapid environmental change. Evolutionary Applications. 2011; DOI:10.1111/j.1752-4571.2010.00166.x

SOS Mata Atlantica. Atlas dos remanescentes florestais da Mata Atlantica: periodo 2000-2005. 2008. http://mapas.sosma.org.br. Accessed 05 Nov 2021.

Swaddle JP, Francis CD, Barber JR, Cooper CB, Kyba CCM, Dominoni DM, et al. A framework to assess evolutionary responses to anthropogenic light and sound. Trends in Ecology & Evolution. 2015; DOI:10.1016/j.tree.2015.06.009

Thatte P, Chandramouli A, Tyagi A, Patel K, Baro P, Chhattani H, et al. Human footprint differentially impacts genetic connectivity of four wide-ranging mammals in a fragmented landscape. Diversity and Distributions. 2020; DOI:10.1111/ddi.13022

Tobler MW. The ecology of the lowland tapir in Madre de Dios, Peru: using new technologies to study large rainforest mammals. Texas A&M University. 2008.

Toews M, Juanes F, Burton AC. Mammal responses to the human footprint vary across species and stressors. Journal of Environmental Management. 2018; DOI:10.1016/j.jenvman.2018.04.009

Tucker MA, Bohning-Gaese K, Fagan WF, Fryxell JM, Moorter BV, et al. Moving in the Anthropocene: global reductions in terrestrial mammalian movements. Science. 2018; DOI:10.1126/science.aam9712

Tucker MA, Busana M, Huijbregts MAJ, Ford AT. Human-induced reduction in mammalian movements impacts seed dispersal in the tropics. Ecography. 2021; DOI:10.1111/ecog.05210

Uezu A, Beyer DD, Metzger JP. Can agroforest woodlots work as steppingstones for birds in the Atlantic Forest region? Biodiversity and Conservation. 2008; DOI:10.1007/s10531-008-9329-0

Varela D, Flesher K, Cartes JL, Bustos S, Chalukian S, Ayala G, et al. Tapirus terrestris. The IUCN Red List of Threatened Species. 2019. https://www.iucnredlist.org/species/21474/45174127. Accessed 05 Nov 2021.

Vaughan D, Dancho M. Furrr: apply mapping functions in parallel using futures. 2021. https://furrr.futureverse.org

Venter O, Brodeur NN, Nemiroff L, Belland B, Dolinsek IJ, Grant, JWA. Threats to endangered species in Canada. BioScience. 2006;56(11):903–910.

Viechtbauer W. Conducting meta-analyses in R with the Metafor Package. Journal of Statistical Software. 2010; DOI:10.18637/jss.v036.i03

Visser AW, Kiorboe T. Plankton motility patterns and encounter rates. Oecologia. 2006; DOI:10.1007/s00442-006-0385-4

Wall J, Wittemyer G, Klinkenberg B, LeMay V, Blake S, Strindberg S, et al. Human footprint and protected areas shape elephant range across Africa. Current Biology. 2021; DOI:10.1016/j.cub.2021.03.042

Wickham H. Ggplot2: elegant graphics for data analysis. Springer-Verlag New York. 2016.

Wood SN. Generalized additive models: an introduction with R. 2nd ed. Chapman; Hall/CRC; 2017.

